# Neural evidence for age-related differences in representational quality and strategic retrieval processes

**DOI:** 10.1101/333484

**Authors:** Alexandra N. Trelle, Richard N. Henson, Jon S. Simons

## Abstract

Mounting behavioral evidence suggests that declines in both representational quality and controlled retrieval processes contribute to episodic memory decline with age. The present study sought neural evidence for age-related change in these factors by measuring neural differentiation during encoding of paired associates, and changes in regional BOLD activity and functional connectivity during retrieval conditions that placed low (intact pairs) and high (recombined pairs) demands on controlled retrieval processes. Pattern similarity analysis revealed age-related declines in the differentiation of stimulus representations at encoding, manifesting as both reduced pattern similarity between closely related events, and increased pattern similarity between distinct events. During retrieval, both groups exhibited increased recruitment of areas within the core recollection network, including the hippocampus and angular gyrus, when endorsing studied pairs, whereas younger adults exhibited increased recruitment of, and hippocampal connectivity with, lateral prefrontal regions during correct rejections of recombined pairs. These results provide evidence for age-related changes in representational quality and in the neural mechanisms supporting memory retrieval under conditions of high, but not low, control demand.

## Introduction

Episodic memory declines in late adulthood; however not all aspects of memory are affected to the same degree. In particular, age-related differences in memory performance are often more pronounced when discriminating between events that share overlapping content, relative to those that are more distinct (Koutstaal & Schacter, 1997; Stark et al., 2013; Devitt & Schacter, 2016). Second, older adults typically exhibit greater memory impairments when demands on recollection-based retrieval processes during retrieval are high, as compared to when memory can be supported by more automatic, familiarity-based processes (Jennings & Jacoby, 1993; Koen & Yonelinas, 2016). This pattern suggests at least two separate factors contribute to age-related memory impairment: a reduction in the availability of high fidelity event representations that can effectively disambiguate between events with overlapping elements; and a decline in the accessibility of these representations, or at least the ability to intentionally retrieve and evaluate these details in a goal-directed manner using controlled retrieval processes. Recent findings from our research group lend support to this two-factor account (Trelle et al., 2017). We found that reducing demands on either the precision of representational content or the recollection-based retrieval processes alone did not eliminate age-related memory deficits, but if both were reduced, age-related differences in memory performance were no longer observed, suggesting a contribution of both of these factors to memory decline in older adults.

Importantly, our work and that of others (Cohn et al., 2008; Luo & Craik 2009) suggests that the ability of older adults to intentionally recall stored details to support memory performance may vary as a function of demands on controlled retrieval processes at test. Such demands can increase as a function of a variety of different task conditions, including the requirement to distinguish between studied targets and novel foils when those foils are experimentally familiar. One paradigm that exemplifies this demand, and which we adopt in the present experiment, is a paired associates recognition memory test. In such a task, participants study a series of paired associates (e.g., word-word pairs or word-picture pairs) and at test are presented with a mixture of studied ‘intact’ pairs and non-studied ‘recombined’ pairs. Critically, these recombined pairs are comprised of studied items, such that one must oppose the familiarity for the individual items in order to correctly assign the recombined pair as non-studied (Lepage et al., 2003; Cohn & Moscovitch, 2007). This is said to necessitate the use of controlled retrieval processes, namely a *recall-to-reject* strategy, which involves the self-initiated elaboration of retrieval cues to deliberately retrieve and evaluate details regarding the correct target associate in order to disqualify the recombined pair as having been studied (Rotello & Heit, 2000; Gallo, 2004). In contrast, the retrieval of target details to endorse intact pairs is said to place considerably smaller demands on such controlled processes at retrieval. In particular, the match between the studied pair and the test cue is said to facilitate access to and retrieval of stored details – or *recall-to-accept* – and so minimize demands on retrieval cue elaboration as well as post-retrieval monitoring and evaluation processes (Rotello & Heit, 2000; Lepage et al., 2003; Cohn & Moscovitch, 2007). Notably, age-related differences in behavioral measures of recall-to-accept are often smaller in magnitude compared to measures of recall-to-reject (Cohn et al., 2008; Trelle et al., 2017). This pattern suggests that instead of a generalized deficit in recollection-based retrieval processes, older adults’ ability to recall stored details may vary according to demands on strategic processes at test.

Collectively, this evidence provides important initial insights into the mechanisms underlying age-related decline in episodic memory, but is inherently limited by the need to use behavioural outcomes (i.e. memory retrieval accuracy) to make inferences about underlying memory representations or the engagement of cognitive control processes. The inability to measure these factors directly makes it difficult to distinguish between deficits that originate during the initial encoding of an event (e.g., reductions in representational specificity), as compared to those that emerge during memory retrieval (e.g., declines in cognitive control processes that support retrieval of stored representations). The present investigation aims to complement existing behavioural findings by using fMRI to identify more direct, neural evidence for each of these constructs. In particular, we use multivariate pattern similarity analysis to measure age-related changes in representational content during memory encoding, and univariate activation and functional connectivity analyses to assess the degree to which age-related differences in retrieval mechanisms vary according to demands on controlled processes at test.

Pattern similarity analysis uses the dissimilarity between voxelwise patterns of neural activity to make inferences about the ability to distinguish different neural representations, such as those of events occurring during memory encoding or retrieval (Kriegeskorte, Mur, & Bandettini, 2008). Using this approach, existing work has provided evidence for the ability to distinguish between visual objects at the level of categories (LaRocque et al., 2013), subcategories (Kriegeskorte et al., 2008), and even individual exemplars (Kriegeskorte, Formisano, Sorger, & Goebel, 2007). These effects tend to be localized in modality-specific areas of sensory cortex, with discrimination of categories often most evident in ventral temporal cortex (Carp et al., 2011; Kriegeskorte et al., 2008). In general, neural activity patterns within sensory cortical regions that are highly responsive to changes in perceptual input are thought to reflect greater neural differentiation, or distinctiveness. That is, greater pattern similarity between related or overlapping events, coupled with lower levels of pattern similarity for unrelated or highly distinct events, reflects greater differentiation (Carp et al., 2011; LaRocque et al., 2013).

Complementing these pattern analysis measures, univariate activation and functional connectivity analyses have been used to examine the recruitment of, and coupling between, brain areas implicated in different aspects of memory retrieval. In particular, although recall-to-accept and recall-to-reject can both result in successful memory retrieval (and potentially equivalent behavioural accuracy), they are known to recruit distinct networks of brain regions in healthy younger adults (Bowman & Dennis, 2017). Successful recollection of a previous event, associated with recall-to-accept, produces increased activity in a set of brain areas, often referred to as the ‘core recollection network’ (Rugg & Vilberg, 2013). These regions include the hippocampus and lateral posterior parietal cortex, particularly within the angular gyrus (Wagner et al., 2005; Vilberg & Rugg, 2008), as well as the medial prefrontal cortex, and retrosplenial/posterior cingulate cortex (Kim, 2010). In contrast, the controlled retrieval processes involved in recall-to-reject produce increased recruitment of dorsolateral prefrontal cortex (DLPFC), particularly BA44 – a region implicated in post-retrieval monitoring and evaluation of retrieved content (Wheeler & Buckner, 2003; Lepage et al., 2003; Achim & Lepage, 2005) – as well as ventrolateral prefrontal cortex (VLPFC), particularly BA47 – a region associated with controlled retrieval of information from long-term memory (Wagner, Maril, Bjork, & Schacter, 2001; Wheeler & Buckner, 2003; Badre &Wagner, 2007). Consistent with this latter proposal, recent work has provided evidence for increased connectivity between the hippocampus and VLPFC during conditions that place increased demands on controlled retrieval processes, such as rejecting familiar lures (Bowman & Dennis, 2017) and retrieving weakly-encoded relative to strongly-encoded source information (Barredo, Oztekin, & Badre, 2015).

The present study combines these measurement tools with a paired associate recognition memory paradigm to examine age-related differences in 1) representational content at encoding and 2) retrieval mechanisms under conditions that vary in their demand on controlled processes. In particular, participants studied word-picture paired associates comprised of trial-unique adjectives paired with one of 8 images (4 objects, 4 scenes), which could be further divided into animate and inanimate objects, and indoor and outdoor scenes. This resulted in word-picture pairs that contained overlapping elements and systematically varied in relatedness to one another, enabling an examination of the degree to which the similarity of stimulus representations during encoding is modulated by perceptual/conceptual overlap, and how this might differ with age. At retrieval, participants were asked to discriminate between previously studied ‘intact’ pairs and non-studied, but experimentally familiar, ‘recombined’ pairs. The pictures from the study phase were removed and replaced with a word label corresponding to the image, eliminating a direct perceptual overlap between studied pairs and their test cues in order to minimize the contribution of automatic processes, such as perceptual fluency, to performance. As such, this design encouraged the retrieval of target details to both accept and reject a given pairing at test. By comparing the neural mechanisms supporting hits to intact pairs and correct rejections to recombined pairs, we could assess the degree to which age-related differences in retrieval processes are modulated by demands on strategic retrieval processes.

We focused our core analyses on a priori regions of interest. Our exploration of representational quality using pattern similarity analysis specifically examined the ventral temporal cortex, comprising parahippocampal gyrus, fusiform gyrus, and lingual gyrus. This broad area has been frequently linked with the representation of visual objects (Carp et al., 2011; Kriegeskorte et al., 2008). To examine the engagement of recollection-based retrieval strategies when endorsing studied items, we selected two key regions of the core recollection network, the hippocampus and angular gyrus, which have each been specifically associated with recollection-based retrieval as compared to recognition based on stimulus familiarity (Eldridge, Knowlton, Furmanski, Bookheimer, & Engel, 2000; Wagner et al., 2005; Yonelinas et al., 2005; Vilberg & Rugg, 2008). The hippocampus plays a critical role in supporting pattern completion processes, or the retrieval of target details in response to a partial cue (O’Reilly & McClelland, 1994), whereas the angular gyrus is said to support the maintenance of retrieved content and the accumulation of mnemonic evidence (Wagner et al., 2005; Vilberg & Rugg, 2008). Finally, to examine the engagement of control processes during correct rejections of recombined items, we focused on the DLPFC and the VLPFC, which support the goal-directed retrieval, selection, maintenance, and monitoring of stored details, particular when control demand is high (Wagner et al., 2001; Wheeler & Buckner, 2003; Badre & Wagner, 2007). We additionally examined functional connectivity between the hippocampus and these cortical regions, which we predicted would vary as a function of strategic retrieval demand (Barredo et al., 2015; Bowman & Dennis, 2017).

During encoding, we predicted that the differentiation of stimulus representations would be reduced in older adults relative to younger adults, and that this would reflect reduced sensitivity to changes in input, manifesting as both reduced pattern similarity between related events, and increased pattern similarity between unrelated events. During retrieval, we predicted that younger adults would exhibit neural patterns consistent with the use of recollection-based retrieval strategies across all trial types. In particular, they should exhibit increased activity in the hippocampus (HIPP) and angular gyrus (ANG) during hits, reflecting target recollection, and increased recruitment of DLPFC, as well as increased hippocampal connectivity with VLPFC, during correct rejections, reflecting the use of recall-to-reject strategy. To the extent that older adults display a generalized recollection deficit, we should observe age-related decline in the presence of each of these neural signatures. In contrast, if older adults’ ability to access stored details to support recognition varies as a function of strategic retrieval demand at test, we should observe an age-invariant neural signature associated with target recollection, that is, increased activity in the hippocampus and angular gyrus during hits relative to correct rejections, coupled with age-related reduction in the mechanisms associated with recall-to-reject, that is, a failure to increase recruitment of DLPFC and VLPFC, and hippocampal-VLPFC connectivity during correct rejections relative to hits.

In summary, the present study sought to characterise age-related differences in both representational quality and retrieval processes under conditions of high versus low control demand, using univariate and multivariate fMRI analysis to directly measuring these constructs during memory encoding and retrieval. In particular, we examined the effects of age on: 1) the differentiation of event representations in ventral temporal cortex during encoding, and how this is modulated by stimulus relatedness, 2) the degree to which the recruitment of “retrieval success” regions (e.g., HIPP, ANG) and “retrieval control” regions (e.g. DLPFC, VLPFC) is modulated by strategic demand during retrieval, and 3) the degree to which HIPP connectivity with these lateral cortical regions (ANG, DLPFC, VLPFC) varies as a function of strategic demand at retrieval.

## Method

### Participants

Twenty younger adults and 22 older adults participated in the study. Participants in both groups were recruited from the MRC Cognition and Brain Sciences Unit volunteer panel as well as the surrounding Cambridge community and received £30 for participating in the study. All participants were healthy, right-handed, had normal or corrected-to-normal vision and hearing, and had no psychiatric or neurological history. Data from two older adults are excluded from the analysis, one due to falling asleep in the scanner and failure to complete the session, and another due to performance below the normal range on the Montreal Cognitive Assessment (MoCA; cut off >= 26; Nasreddine et al., 2005). All remaining older adults performed within the normal range on the MoCA (*M* = 27.8). Older (*M* = 71.4 years; 8F, 12M) and younger (*M* = 24.9 years; 13F, 7M) adults did not differ with respect to years of formal education (*t*(38) = 1.60, *p* = .118), and older adults performed higher on the Shipley Institute of Living Scale (*t*(38) = 3.55, *p* < .001). Informed consent was obtained in accordance with the Cambridge Psychology Research Ethics Committee.

### Materials

Experimental stimuli were 192 word-picture pairs comprised of trial-unique adjectives paired with one of eight colored pictures. Four of these pictures were of objects, two of which were living things and two were inanimate objects, and the remaining four pictures were of scenes, two depicting indoor settings and two outdoor settings (see Figure 1). Thus, the resulting paired associates systematically varied in relatedness. That is, some pairs shared a common stimulus category and overlapping content (e.g. MUDDY Umbrella and GOLDEN Umbrella), others shared a common subcategory, but non-overlapping content, (e.g. MUDDY Umbrella and WOODEN Teapot), others shared a common category but different subcategory (e.g. MUDDY Umbrella and SPOTTED Rabbit), and others contained non-overlapping content from the opposite stimulus category (e.g. MUDDY Umbrella and STRIPED Office). Adjective-picture pairings were fixed across participants, and were designed to ensure that the picture could plausibly be imagined in accordance with the adjective.

**Figure 1.**
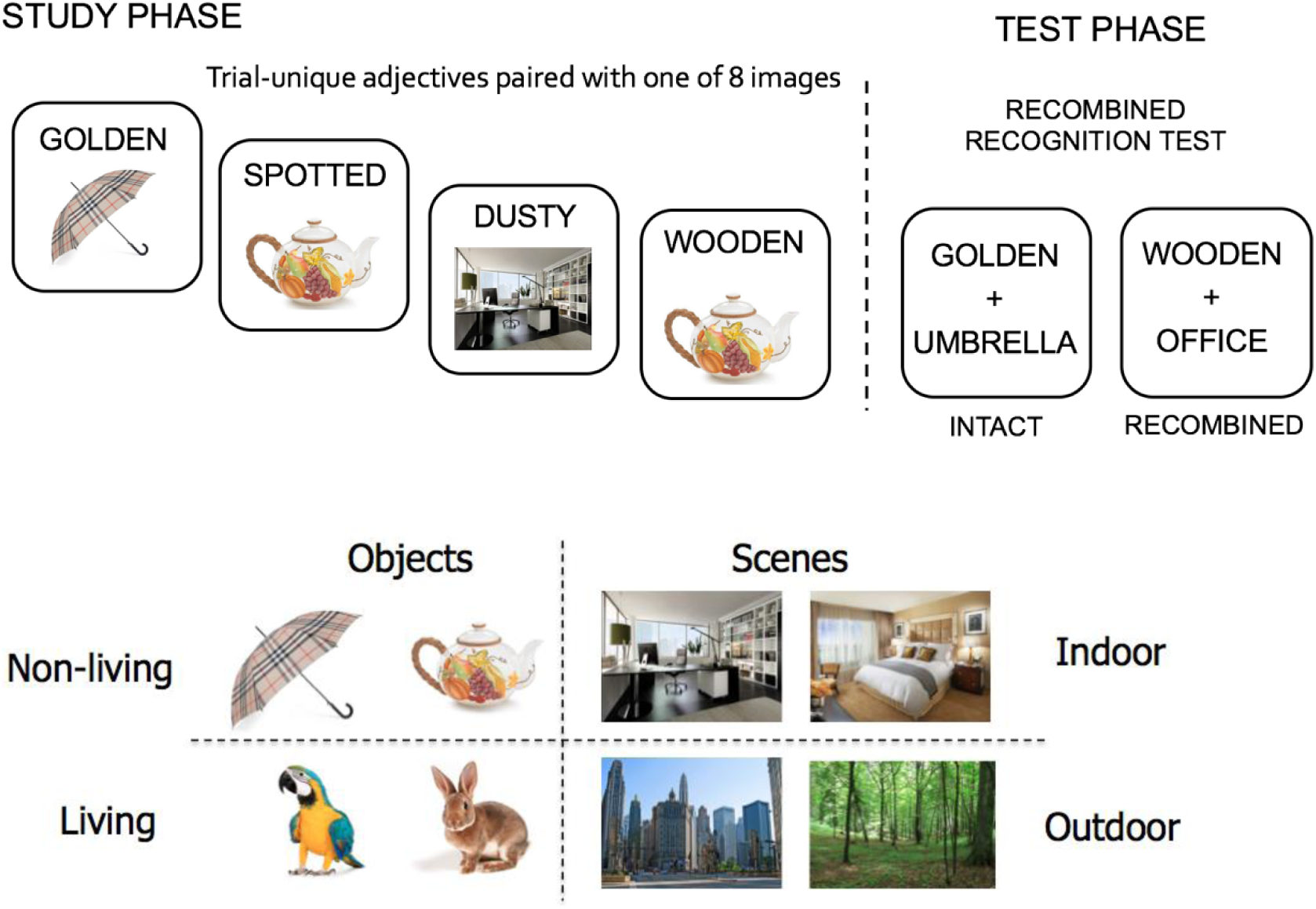
Schematic depicting experimental paradigm. Participants studied trial-unique adjectives paired with one of eight images pictured above. At test, participants were presented with studied (intact) and non-studied (recombined) pairs and made an old/new judgment for each. Pictures were replaced with word labels during the test phase to reduce perceptual overlap between study and test, minimizing the ability to rely on perceptual fluency to support performance.

Word-picture pairs were randomly assigned to one of three 64-item study lists, with the constraint that each list contained eight pairs corresponding to each of the eight pictures. Half of the study items were subsequently presented as intact pairs during the test phase, and the other half were presented as recombined pairs. The assignment of pairs as intact or recombined was counterbalanced across participants. During the test phase, each associate image was replaced with a common noun denoting the image (e.g., TEAPOT, BEDROOM), to eliminate direct perceptual overlap between study and test trials for intact pairs and minimize the ability to rely solely on perceptual fluency to endorse intact pairs. To ensure that participants understood which label corresponded to which picture, the word labels were described to participants prior to beginning the task. The presentation of word-picture pairs during each study and test block was pseudo-randomized for each participant, with the constraint that no more than four images from the same category appeared in sequence, and that each image was not presented more than twice in a row. The test phase included the additional constraint that no more than four intact or recombined pairs occurred in sequence. Stimuli were presented using the Cogent software package implemented in MATLAB (Mathworks, Inc., USA).

### Procedure

The experimental paradigm is depicted in Figure 1. Each study block comprised 64 trials in which participants were presented with a word-picture pair and instructed to imagine the picture in accordance with the adjective (e.g., to imagine a golden umbrella), and to indicate whether they had been successful in doing so with a button press response (1 = successful, 2 = unsuccessful). Each study block was followed by a one-minute retention interval during which, to prevent rehearsal, participants were asked to covertly count backwards from a random number presented on the screen. The test phase commenced immediately afterwards. During each test trial, participants were presented with a studied adjective and a word corresponding to one of the eight pictures and asked to indicate with a single button press whether each pairing was previously studied together (old) or recombined (new). Both study and test trials lasted for a fixed duration of 5000 ms, with an inter-trial interval of 1000 ms. Responses not made within the allotted time were marked as no response and excluded from the subsequent analysis. Participants completed three alternating study-test blocks in this fashion, with a one-minute break between each cycle during which they were instructed to close their eyes and rest. The Shipley Vocabulary test and Montreal Cognitive Assessment were completed at the end of the session outside of the scanner.

### fMRI Data Acquisition and Pre-processing

Scanning was performed using a 3-T Siemens Prisma MRI system with a 32-channel head coil. Functional data was acquired using a descending Blood-Oxygenation-Level-Dependent (BOLD)- weighted echo-planar imaging (EPI) pulse sequence (repetition time (TR) = 2000 ms, echo time (TE) = 30 ms, flip angle = 78). Each EPI volume consisted of 32 axial slices (3mm thick, 0.75 mm gap, 3 × 3 mm in-plane resolution) covering the whole brain. For each of the six sessions (3 study and 3 test blocks), 210 volumes were acquired. The first five volumes of each session were discarded to allow for magnetic field stabilization. A high-resolution (1 × 1 × 1 mm) T1-weighted anatomical image was also acquired at the beginning of the scanning session using a 3D magnetization-prepared rapid acquisition gradient echo (MP-RAGE) pulse sequence.

Data pre-processing and univariate analysis was conducted using SPM12 (http://www.fil.ion.ucl.ac.uk/spm/) and batched using “automatic analysis” software (version 4; https://github.com/rhodricusack/automaticanalysis/). Preprocessing of image volumes included spatial realignment to correct for movement, followed by slice-timing correction, using the first acquired slice in each volume as a reference. The structural image of every participant was registered to a sample-specific template using the diffeomorphic flow-field method of DARTEL (Ashburner, 2007), and the template subsequently transformed to the Montreal Neurological Institute (MNI) stereotactic space using a 12-parameter affine transformation. The mean functional volume of each participant was coregistered to their structural image, and the DARTEL plus affine transformations applied to transform the functional images into MNI space.

### Regions of Interest

Analyses examining age-related differences in the differentiation of stimulus representations during encoding focused on ventral temporal cortex. This region was functionally defined using independent data from the same participants obtained during a localizer run, in which participants viewed all eight images from the main experiment, with each of the eight images presented 12 times for a total of 96 events (duration = 4000ms, ISI = 1000ms). During the localizer run, participants performed a simple colour detection task, in which they indicated whether a small square placed in a random location on the screen was either pink or yellow, to ensure attention to the stimuli. A whole-brain, Group (Young, Old) × Stimulus Type (Object, Scene) ANOVA was conducted to identify object- and scene-selective cortex common to both groups. The results of the contrasts Objects > Scenes and Scenes > Objects were thresholded at *p* < .05 FWE, and masked to include only anatomical regions within bilateral ventral temporal cortex, including parahippocampal cortex, fusiform gyrus, and inferior temporal cortex, as defined by the automated anatomical labelling (AAL) atlas (see Figure 2).

**Figure 2:**
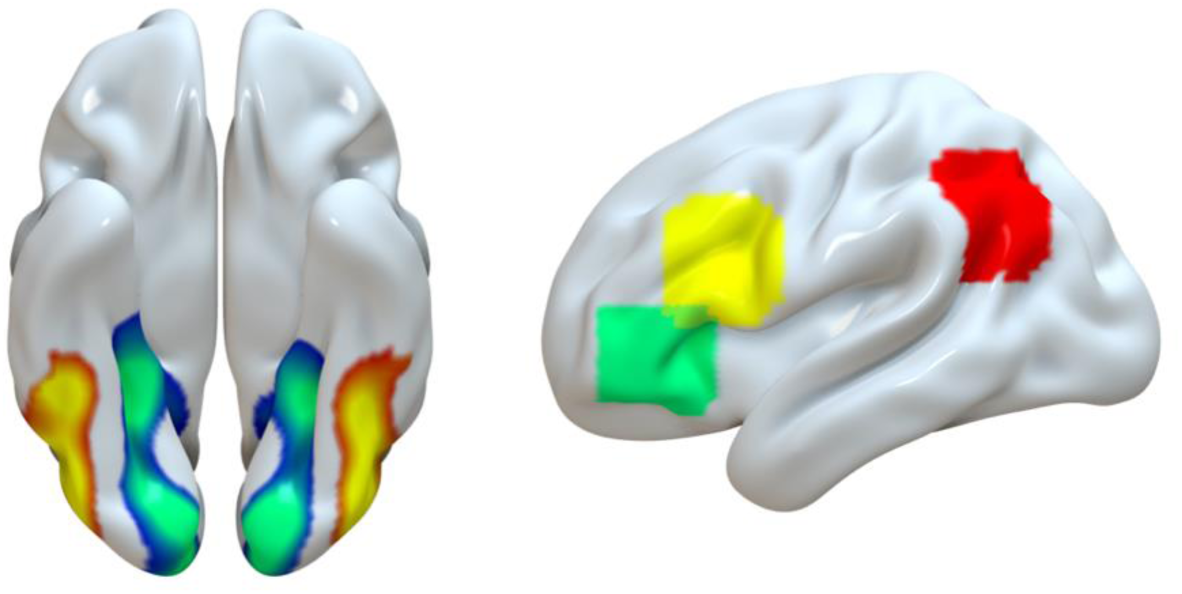
Cortical Regions of Interest (ROIs). Left panel: Age-invariant results of the contrast Objects > Scenes (yellow-red) and Scenes > Objects (green-blue) contrasts during the localizer task, thresholded at p < .05 FWE corrected, used as a mask for analysis of representational quality during encoding. Right panel: Coordinate-based ROIs defined by Wheeler & Buckner (2003) centered on a peak coordinate in DLPFC (yellow; peak coordinates −47,+17,+24), VLPFC (green; peak coordinates −45, +35, −4), and Angular Gyrus (red; peak coordinates −45, −69, −6).

Univariate analysis of age-related differences in retrieval processes focused on four a priori anatomical regions of interest (ROIs) within the left hemisphere. Two areas associated with retrieval success, including the hippocampus (HIPP), the angular gyrus (ANG), and two regions associated with retrieval control, namely ventrolateral prefrontal cortex (VLPFC), approximating BA47, and dorsolateral prefrontal cortex (DLPFC), approximating BA44. The ANG, VLPFC, and DLPFC masks were defined as 10mm spheres centered on peak coordinates from a previous investigation examining the contributions of these regions to perceived oldness and cognitive control during retrieval, respectively (Wheeler & Buckner, 2003; see Figure 2). The spatial localization of these coordinates within each anatomical region corresponds well with the coordinates reported in previous work examining the role of these regions during memory retrieval (Badre & Wagner, 2007; Lepage et al., 2003; Achim & Lepage, 2005; Wagner et al., 2005; Vilberg & Rugg, 2008; Barredo et al., 2015; Bowman & Dennis, 2017). The HIPP was defined anatomically based on the automated anatomical labelling (AAL) atlas.

Functional connectivity analyses examining age-related differences in retrieval processes were conducted within the same set of a priori ROIs, but this time focusing on connectivity between the HIPP and the remaining cortical regions associated with retrieval, namely ANG, VLPFC, and DLPFC. The HIPP seed region was defined in a participant-specific manner as a sphere of 5mm radius centered on each individual’s peak BOLD activity within the hippocampus during retrieval, identified using the first-level contrast of Hits > CRs, in order identify the precise area within the hippocampus that was engaged during the task for each individual.

### Univariate Analysis

Univariate analyses were used to examined differences in regional BOLD activity during retrieval under conditions of high (correct rejections) and low (hits) retrieval control demand. For this analysis, EPI images were smoothed with an isotropic 8-mm full-width-at half-maximum (FWHM) Gaussian kernel before modelling. Neural activity was modeled by delta functions at stimulus onset for each event of interest and convolved with a canonical hemodynamic response function (HRF). The resulting timecourses were downsampled at the reference slice for each scan to form regressors in the General Linear Model (GLM). The model contained two regressors of interest representing the two types of successful retrieval events: hits and correct rejections. All remaining trials (misses, false alarms, no response) formed a third regressor of no interest, and 6 additional regressors representing movement parameters estimated during spatial alignment (3 rigid-body translations, 3 rotations) were additionally included. Voxel-wise parameter estimates for each regressor were obtained by restricted maximum-likelihood estimation, using a temporal high pass filter (cut-off 128 s) to remove low-frequency drifts and an AR(1) model of temporal autocorrelation.

We first examined changes in regional BOLD activity in four a priori regions of interest (see *Regions of Interest)* associated with retrieval success and retrieval control. Specifically, we extracted the first-level parameter estimates (scaled to represent percent signal change) from each region for each participant, and submitted these values to a mixed ANOVA with factors Group (Young, Old) and Trial Type (Hit, CR). The outcome of this test for each ROI was considered significant if the p value survived Bonferroni correction for multiple comparisons (.05/4 regions = .0125).

We additionally conducted a whole-brain voxelwise analysis to identify areas outside of our a priori ROIs that were differentially recruited during hits and correct rejections, and the degree to which this differed with age. First-level contrasts of the parameter estimates (Hit, CR) for each participant were entered into a second-level Group (Young, Old) and Trial Type (Hit, CR) mixed ANOVA examine both main effects and condition-by-group interactions. All effects were thresholded at a whole-brain significance level of *p* < .05 FWE corrected. Whole-brain maps applying a more lenient threshold of *p* < .005 uncorrected with a minimum cluster size of 10 contiguous voxels are available in the supplementary materials. Note that we did not examine retrieval success effects (i.e. differences in regional BOLD activity during remembered vs forgotten trials) due to the low frequency of incorrect trials in the majority of the younger participants.

### Functional Connectivity Analysis

Functional connectivity analyses examined hippocampal-cortical connectivity under conditions of high (correctly rejecting recombined pairs, CRs) relative to low (correctly endorsing intact pairs, Hits) retrieval control demand. Beta estimates for each test trial were obtained by modelling smoothed EPI data from all three test sessions in a single GLM containing a separate regressor for each trial, as well as six movement parameters and the mean for each session as regressors of no interest. This procedure yielded separate beta images corresponding to each of the 192 test trials, which were submitted to functional connectivity analyses using the beta-series correlation method (Rissman, Gazzaley, & D’Esposito, 2004), which quantifies functional connectivity as the Pearson correlation between the beta-series. In particular, we conducted a whole brain seed-to-voxel connectivity analysis during hits and correct rejections, using a participant-specific hippocampal ROI as a seed region (see *Regions of Interest*). The hippocampal beta series calculated as the mean beta series of all voxels (19) within the 5mm sphere.

We first examined hippocampal connectivity within three a priori target regions of interest, namely the three remaining cortical regions of interest (i.e., ANG, VLPFC, DLPFC; see Figure 2) used for the univariate analyses and described in *Regions of Interest*. Specifically, we averaged the timeseries within each target ROI and computed the mean correlation between the hippocampus and each target region for each trial type. We then submitted these values to a Group (Younger, Older) x Trial Type (Hit, CR) mixed ANOVA to explore main effects of retrieval control demand on connectivity, and how this might differ across age groups. To correct for multiple comparisons across the three regions of interest, only those p values surviving correction for multiple comparisons (.05/3 = 0.0167) were considered significant. As for the univariate analyses described above, we additionally examined the results of the whole-brain seed-to-voxel connectivity analysis, in which the first-level participant-specific beta-series correlation maps were entered into a second-level Group x Trial Type ANOVA. No effects exceeded a threshold of *p* < .05 with FWE correction. Whole brain maps using a more lenient threshold (*p* < .005 uncorrected with a minimum cluster size of 10 voxels) can be found in the supplementary materials.

### Representational Similarity Analysis

Representational similarity analysis (Kriegeskorte et al., 2008) was used to examine the differentiation of stimulus representations during encoding in ventral temporal cortex (VTC; see *Regions of Interest for definition*). Beta estimates for each study trial were obtained by modelling unsmoothed data from all three study sessions in a single GLM containing a separate regressor for each trial, as well as six movement parameters and the mean for each session as regressors of no interest. This procedure yielded separate beta images corresponding to each of the 192 study trials. We computed the Pearson correlation between all encoding trials, irrespective of subsequent memory accuracy (see Supplementary Data for results using only correct trials, which did not affect the pattern of results), producing a 192 × 192 correlation matrix for each participant. These correlation values were Fisher transformed before computing the mean pairwise correlation between events corresponding to each of four event types, using across-run pairs only to compute mean pattern similarity for each conditions. These event types were defined based on the image exemplar presented on each trial: Same Exemplar (e.g. MUDDY Umbrella and GOLDEN Umbrella), Same Subcategory (e.g. MUDDY Umbrella and WOODEN Teapot), Same Category (e.g. MUDDY Umbrella and PAINTED Rabbit) and Different Category (e.g. MUDDY Umbrella and STRIPED Office). Here we report the mean pairwise correlation for each event type and conduct a Group x Level ANOVA to examine the effect of stimulus relatedness on pattern similarity across age groups.

We additionally compute the difference scores between successive levels of relatedness (SE-SS, SS-SC, SC-DC) to estimate exemplar, subcategory, and category distinctiveness (Carp et al., 2011), which reflect the magnitude of the difference in pattern similarity between successive levels of event similarity. Difference scores significantly greater than zero as assessed using an independent samples t-test provide evidence for discriminability of a given level of granularity in neural activity patterns in ventral temporal cortex.

## Results

### Behavioural Results

Participants’ responses were categorized as hits, correct rejections, false alarms, or misses. Trials in which participants did not respond within the allotted 5 seconds were marked as ‘no response’, and made up 1% and 2.5% of trials in younger and older adults, respectively. The number of trials comprising each trial type is presented in Table 1. Recognition memory performance was calculated as the proportion of hits to intact pairs corrected by the proportion of false alarms to recombined pairs. Independent samples *t-*tests revealed that memory performance was significantly impaired in older adults relative to younger adults, with older adults making fewer hits (*t*(38) = 4.67, *p* < .001, *d* = 1.52) and fewer correct rejections (*t*(38) = 5.75, *p* < .001, *d* = 1.87) relative to younger adults (see Figure 3). Due to the low frequency of incorrect trials among the majority of the younger adults (misses: *M* =9.95, *SD* = 6.6; false alarms: *M* =5.75, *SD* = 4.2), we restrict our primary analyses to hits and correct rejections.

**Figure 3:**
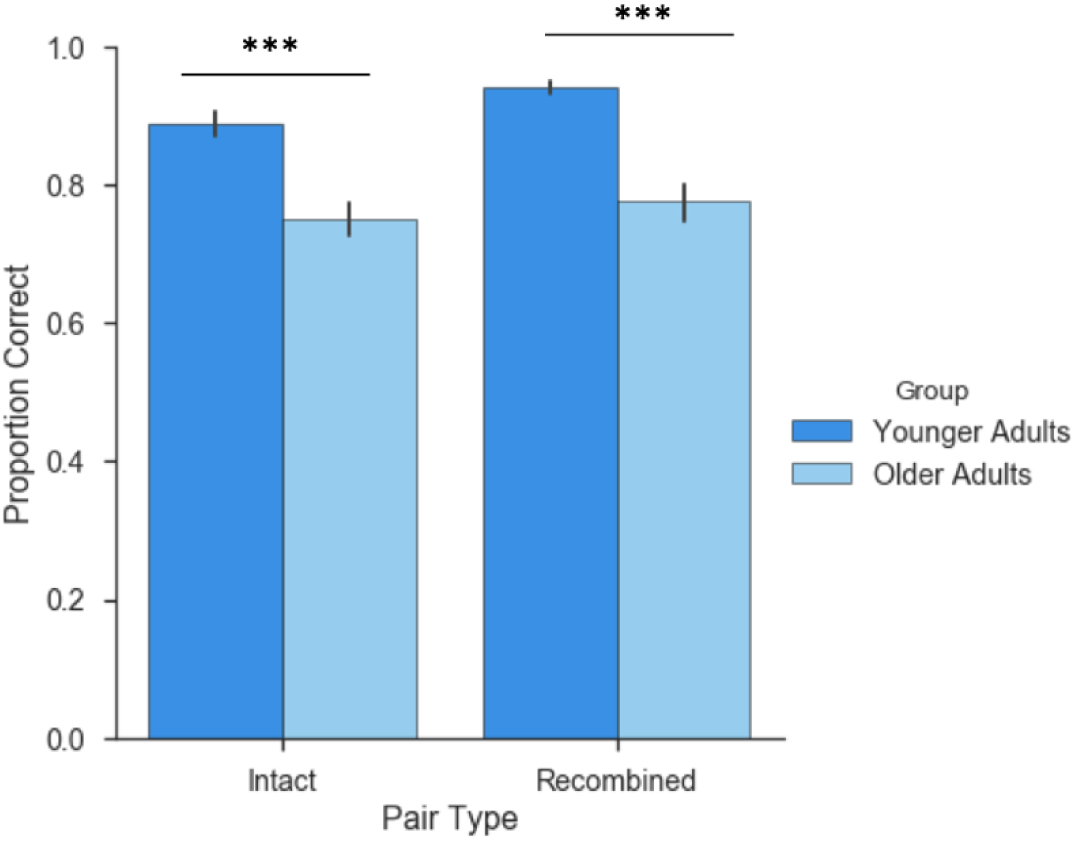
Behavioural Performance. Mean proportion of hits to intact pairs and correct rejections of recombined pairs. Error bars represent standard error of the mean. Older adults made fewer hits and correct rejections relative to younger adults, *** p < .001.

**Table 1:** Mean (SD) trial counts for each response type by age group

|  | Hits | CRs | FAs | Misses |
| --- | --- | --- | --- | --- |
| Younger Adults | 85.05 (7.3) | 89.60 (4.8) | 5.75 (4.2) | 9.95 (6.6) |
| Older Adults | 71.90 (10.3) | 72.0 (11.8) | 21.40 (11.5) | 21.95 (9.2) |

### Stimulus Differentiation in VTC during Encoding

To assess age-related change in the differentiation of representational content, we tested whether pattern similarity between VTC stimulus representations at encoding varied as a function of event type (SE, SS, SC, and DC), and whether this differed with age (Figure 4). A 2×4 ANOVA revealed a main effect of event-type, *F*(3,151)=24.17, *p* < .001, with greatest pattern similarity between same exemplars (SE) and least between exemplars from the different category (DC). More importantly, though the main effect of age group was not significant (*F*<1), there was a significant interaction between event-type and age group (*F*(3,151)=8.94, *p* < .001). Follow-up t-tests revealed that the older group exhibited lower pattern similarity between events sharing the same exemplars (*t*(38)=2.37, *p* =.023) and events sharing the same subcategory (*t*(38)=2.22, *p* =.033) than the younger group, but greater pattern similarity between exemplars of the different category relative to the younger group (*t*(38)=4.45, *p* <.001). Events from within the same category did not differ (*t*(38)=1.23, *p* =.227). Thus, older adults exhibit both decreased pattern similarity for perceptually and conceptually similar events, as well as increased pattern similarity for events that are more distinct, yielding a net decline in discriminability.

**Figure 4:**
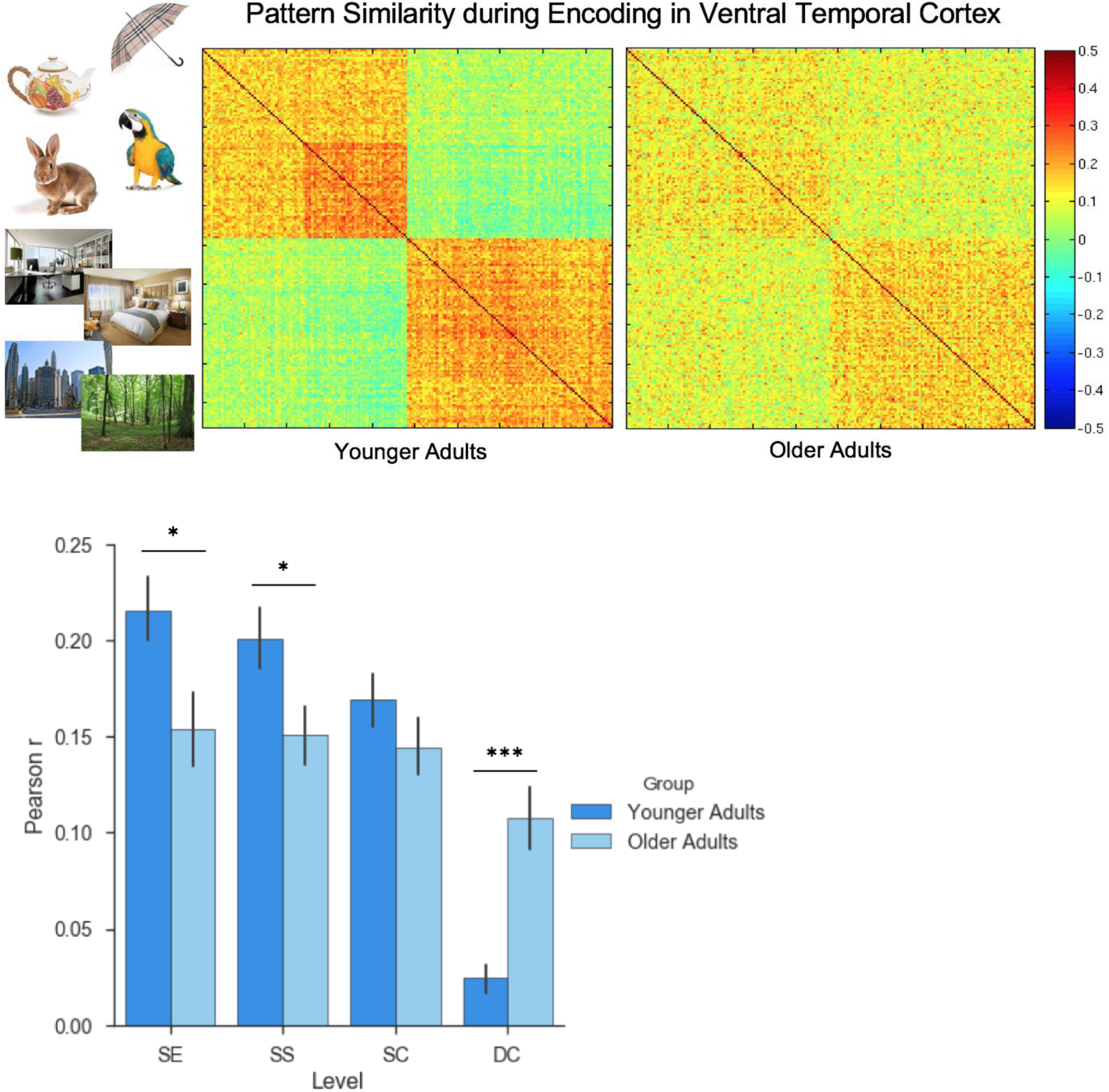
Pattern similarity at encoding as a function of event relatedness. Top: Correlation matrix depicting the Pearson correlation between individual encoding events, sorted by category, subcategory, and then by the exemplar presented on each trial. Bottom: Mean correlation between events as a function of event relatedness. Relative to younger adults, older adults exhibit reduced pattern similarity for related events, coupled with increased pattern similarity for events that are more distinct. Error bars represent standard error of the mean. ***p < .001; * p < .05

**Figure 5:**
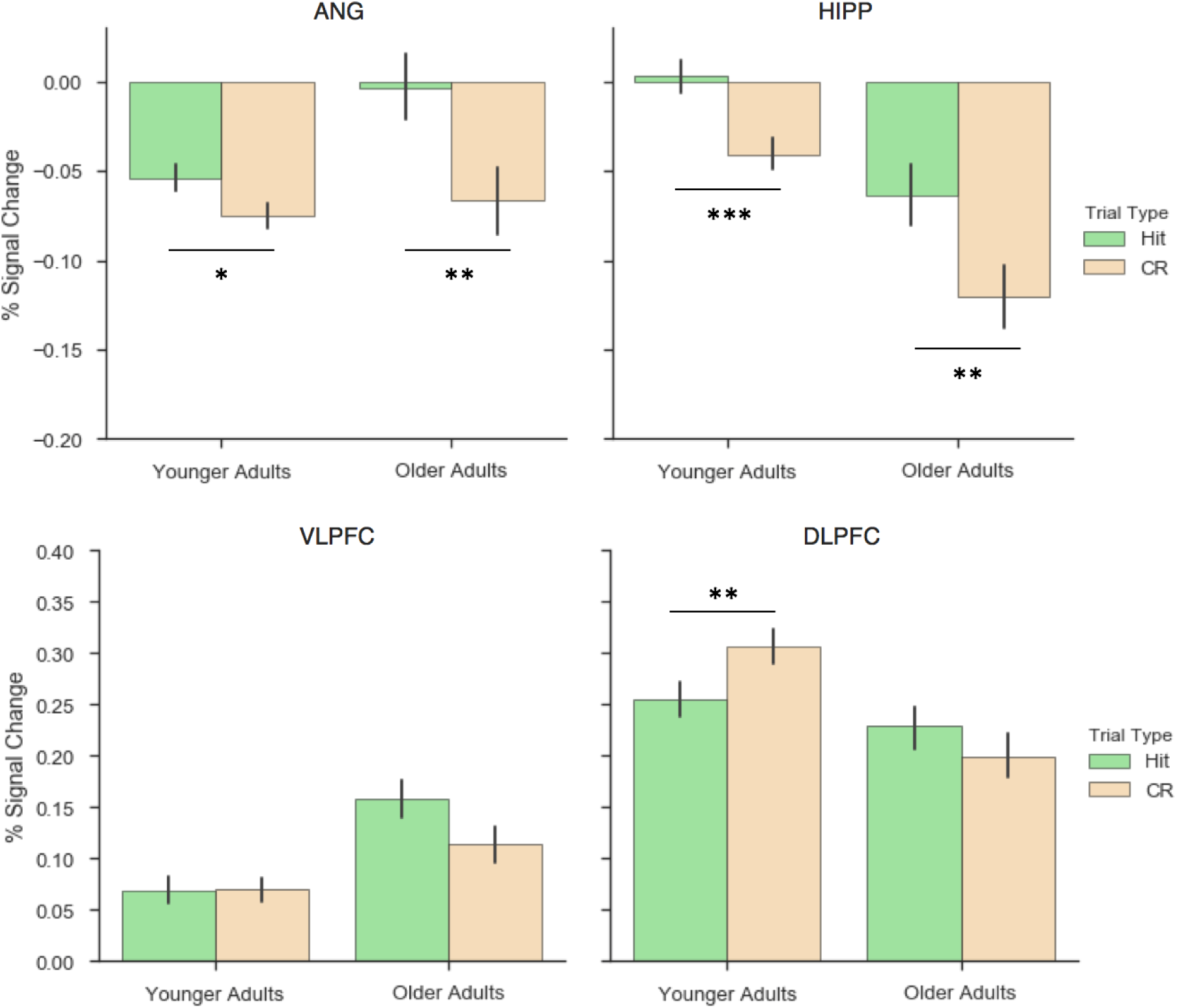
Percent signal change during Hits and CRs in each regions of interest. Both older and younger adults exhibited increased activity in ANG and HIPP during Hits relative to CRs. Only younger adults exhibited increased activity in DLPFC during CRs relative to Hits. Error bars represent standard error of the mean. ***p < .001; ** p < .01; * p < .05

To measure exemplar, subcategory, and category discriminability, we computed difference scores between subsequent levels of the stimulus hierarchy (e.g., SE-SS, SS-SC, SC-DC), where difference scores significantly greater than zero indicate that a given level can be discriminated on the basis of activity patterns in ventral temporal cortex. Among younger adults, exemplars (*t*(19) = 2.50, *p* = .022), subcategories (*t*(19) = 7.36, *p* < .001), and categories (*t*(19) = 8.97, *p* < .001) were each discriminable, whereas among older adults this was only true for stimulus category (*t*(19) = 6.15, *p* < .001; exemplars: *t* <1; subcategories: *t*(19) = 1.19, *p* = .247).

### Changes in Regional Univariate BOLD Activity during hits and correct rejections

To assess age-related differences in the engagement of strategic processes during retrieval, we next examined the recruitment of ‘retrieval success’ regions and ‘retrieval control’ regions during hits and correct rejections. To this end, we conducted a 2 (Trial Type) × 2 (Group) mixed ANOVA over the parameter estimates from each participant’s first-level models for hits and correct rejections in each ROI. Effects were considered significant if the p-value fell below the Bonferroni corrected threshold *p* < .0125. In HIPP, this revealed a main effect of Trial Type (*F* (1,38)=24.49, *p* < .001), but no effect of Group (*F*(1,38)=1.12, *p* =.297) and no Group x Trial Type interaction (*F* < 1), with both groups exhibiting increased hippocampal activity during hits relative to CRs. Similarly, in ANG there was a main effect of Trial Type (*F* (1,38)=16.60, *p* < .001), with greater activity for hits than correct rejections, but no difference between Groups (*F* < 1) nor a Group x Trial Type interaction (*F* (1,38)=4.17, *p* =.048). In DLPFC, there was a Group x Trial Type interaction (*F* (1,38)=8.12, *p* < .007), but no main effect of Group (*F* < 1) or Trial Type (*F* < 1), reflecting an increase in DLPFC activity for CRs relative to hits in younger (*t*(19) = 2.98, *p* < .007), whereas older adults show a numerical but non-significant difference in the opposite direction (*t*(19) = 1.3, *p* =.206). In VLPFC, main effects of Group (*F* <1) and Trial Type (*F*(1,38)=3.47, *p* = .07) were not significant, nor was the Group x Trial Type interaction (*F*(1,38)=3.96, *p* = .05).

### Changes in Functional Connectivity during hits and correct rejections

To further explore age-related differences in the ability to engage strategic retrieval processes, we next assessed whether hippocampal connectivity with each lateral cortical region of interest varied as a function of demands on strategic retrieval, and whether this differed as a function of age (Figure 6). To this end, we conducted a 2 (Trial Type) × 2 (Group) mixed ANOVA for each ROI over the mean hippocampal-cortical correlation for each trial type. Effects were considered significant if the p-value fell below the Bonferroni corrected threshold of *p* < .0167. In VLPFC, this revealed a Group x Trial Type interaction (*F* (1,38)=9.28, *p* < .005), reflecting an increase in HIPP-VLPFC connectivity during CRs relative to hits in younger (*t* (19) = 4.03, *p* < .001), but not older adults (*t* (19) = .64, *p* = .525). The similar qualitative pattern emerged in DLPFC, but here the Group x Trial Type interaction was not significant (*F* (1,38)=4.05, *p* =.051). However, a main effect of Group (*F* (1,38)=7.35, *p* <.01) was observed, indicating greater HIPP-DLPFC connectivity in older relative to younger adults overall. In ANG, the Group x Trial Type interaction was not significant (*F* (1,38)=2.05, *p* =.16), nor was the main effect of Trial Type (*F* (1,38)=1.63, *p* =.21), but a main effect of Group emerged (*F* (1,38)=6.92, *p* =.0123) again indicating greater connectivity among older adults than younger adults that did not vary with trial type.

**Figure 6:**
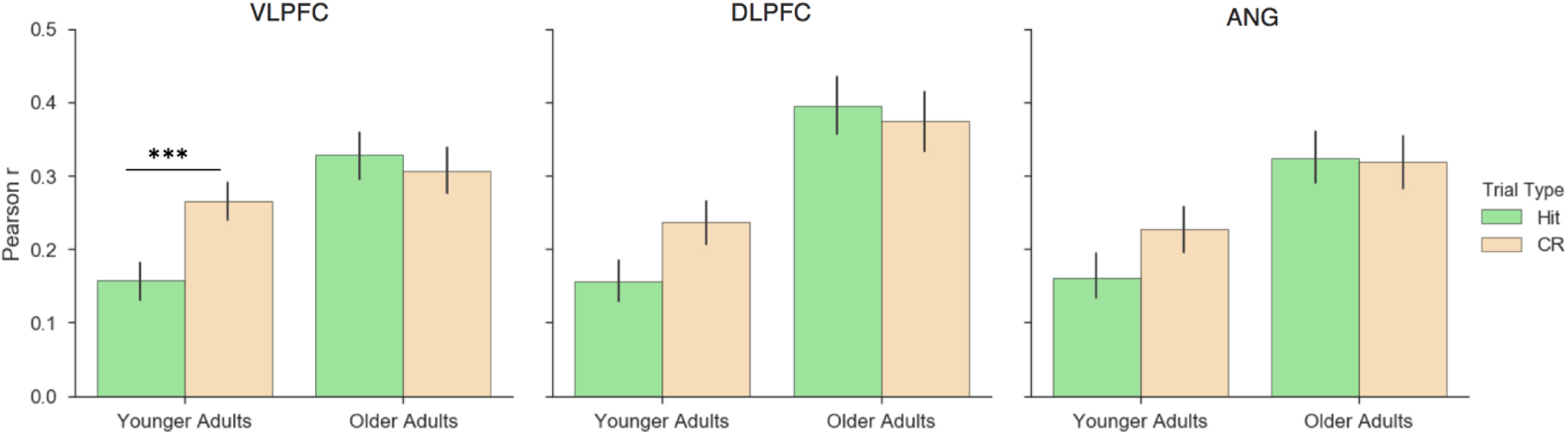
Mean hippocampal connectivity (Pearson r) with each cortical region of interest during Hits and CRs. Only younger adults exhibited increased hippocampal connectivity with VLPFC during CRs relative to Hits, whereas hippocampal connectivity did not vary across trial types in older adults. Error bars represent standard error of the mean. ***p < .001

**Figure 7:**
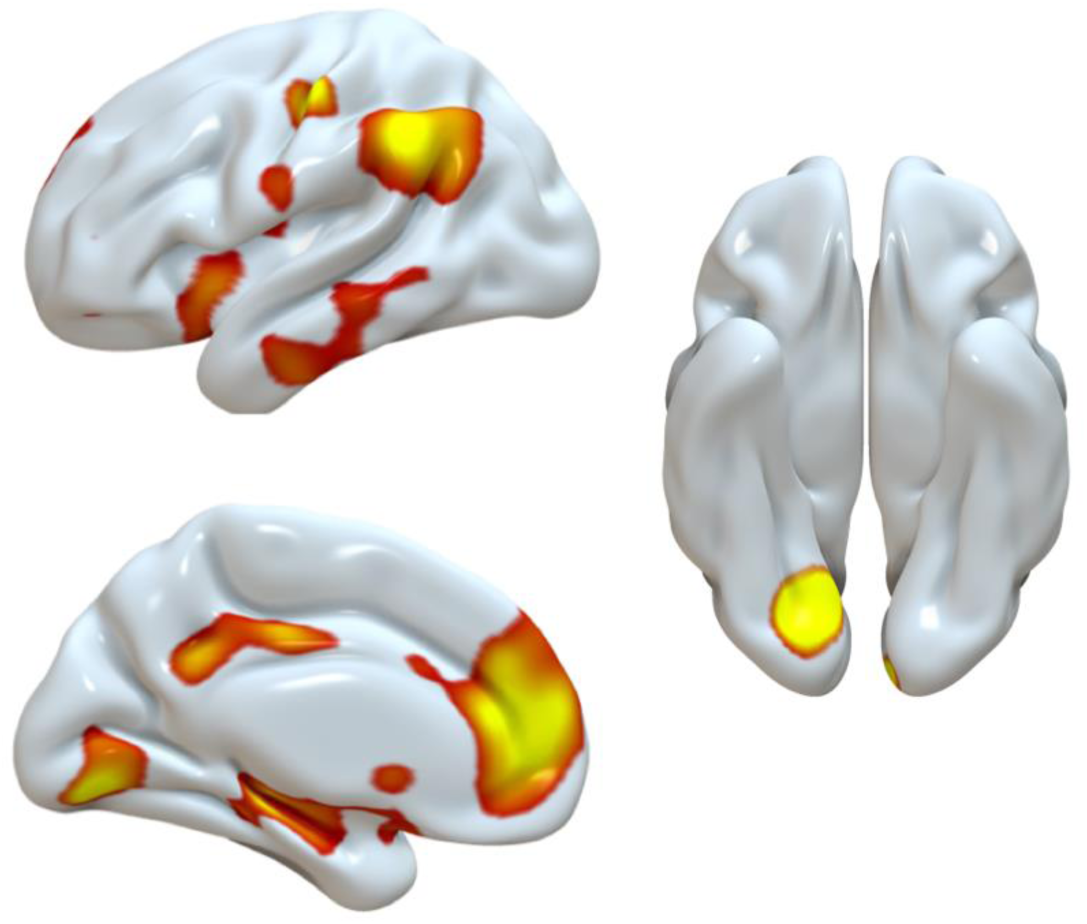
Age-invariant univariate effects during Hits > CRs (left two panels) and CRs > Hits (right panel), p<.05 FWE corrected.

### Whole-brain Exploratory Analysis

To identify areas differentially engaged during hits and correct rejections across age groups beyond the a priori ROIs described above, we next conducted a whole-brain voxelwise univariate analysis (see Table S2). The contrast of Hit>CR revealed an age-invariant increase in activity in brain areas within the core recollection network (Rugg & Vilberg, 2013), including the hippocampus, angular gyrus, medial prefrontal cortex, and posterior cingulate cortex, as well as regions within ventral visual and occipital cortex. The CR>Hit contrast yielded one cluster in lingual gyrus. The Age x Condition interaction did not yield any significant clusters at *p* < .05 FWE (but see Supplemental Data for whole brain uncorrected maps). Whole brain seed-voxel connectivity analysis using the same participant-specific hippocampal seed regions used in the ROI analysis described previously yielded no main effects wherein hippocampal connectivity differed between hits or correct rejections at a threshold of *p* < .05 FWE, nor any age x condition interactions (but see Supplementary Data for whole brain uncorrected maps).

## Discussion

The present study used univariate and multivariate analyses of fMRI data to investigate potential age-related changes in neural differentiation during memory encoding and in the engagement of strategic retrieval processes during memory retrieval. In particular, we measured neural differentiation during encoding of paired associates that systematically varied in perceptual and conceptual relatedness, and changes in regional BOLD activity and hippocampal connectivity during retrieval conditions that placed low (intact pairs) and high (recombined pairs) demands on strategic retrieval processes. We obtained evidence for age-related reductions in the differentiation of stimulus representations during encoding, as well as age-related changes in the neural mechanisms associated with rejecting recombined pairs (likely to reflect recall-to-reject strategies), coupled with age-invariant mechanisms for endorsing intact pairs (likely to reflect recall-to-accept strategies). These results suggest that age affects both representational quality and retrieval control processes, consistent with existing behavioural evidence (Trelle et al, 2017), but using fMRI to more directly measure these constructs, and provide insights into the neural mechanisms that support successful memory retrieval in older adults.

Consistent with hypotheses, we observed an age-related reduction in the differentiation of neural representations for stimuli during encoding. In particular, whereas neural activity patterns in ventral temporal cortex could distinguish between events at the level of categories, subcategories, and exemplars in younger adults, only events from different stimulus categories (i.e., objects, scenes) could be reliably discriminated within the older adult data. This age-related reduction in the discriminability of events during encoding manifested as both a reduction in pattern similarity for highly similar events (i.e. those sharing the same exemplar) coupled with increased pattern similarity for highly distinct events (i.e. those from the opposite stimulus category), relative to younger adults. This pattern suggests that reductions in representational quality with age are driven by a decline in both the *sensitivity* of neural responses to different stimuli and the *selectivity* of neural responses to different stimuli. This reduction in the differentiation of event representations may contribute to declines in the availability of high precision details during memory retrieval, leading to age-related impairments in the ability to discriminate between events that are highly similar or share overlapping elements, relative to those that are more distinct.

These results complement existing findings identifying reductions in the distinctiveness of visual category representations with age (Park et al., 2004; Carp et al., 2011), and extend this work by examining not only category-level differentiation, but also discriminability of subcategories and individual exemplars. Importantly, we did not identify evidence for discriminability of event representations at these more fine-grained levels in older adults, highlighting the importance of not only examining neural differentiation at the categorical level, which is common in experimental aging studies, but also differentiation at finer scales, which may more closely approximate the discriminations one faces in daily memory decisions. The present findings diverge however from some previous observations of age-invariant pattern discrimination between stimuli during encoding, such as audiovisual clips (St-Laurent et al., 2014). This difference across studies might reflect the benefit to stimulus representations afforded when the nature of the information is more complex, e.g., multi-modal, thus automatically providing a more differentiated input and facilitating the differentiation of neural representations. Taken together, these data suggest that although dedifferentiation may occur with age, there are steps that can be taken to enhance representational quality, including the nature of the stimuli and encoding operations.

In addition to declines in representational quality during encoding, we predicted that age-related differences would be observed during memory retrieval, particularly when demands on strategic retrieval processes are high (i.e. when rejecting recombined pairs). Consistent with this hypothesis, we observed an increase in the recruitment of DLPFC during correct rejections relative to hits in younger adults but not older adults. This pattern of increased recruitment replicates previous work in younger adults, and has been linked to increased post-retrieval monitoring demands associated with disqualifying a recombined pair as having been studied (Lepage et al., 2003; Achim & Lepage, 2005). Similarly, only younger adults exhibited increased coupling between the hippocampus and VLPFC during correct rejections relative to hits, a pattern of connectivity that has previously been linked to the goal-directed retrieval and selection of target details during the execution of a recall-to-reject strategy (Barredo et al., 2015; Bowman & Dennis, 2017). Instead, older adults exhibited greater overall connectivity between the hippocampus and DLPFC and ANG, but this did not vary by trial type. As it is possible that this overall increase in connectivity simply reflects age-related differences in baseline connectivity between brain areas (Blum et al., 2014; Ferreira et al., 2016), here we focus on the differences, or lack thereof, between conditions. The observation that older adults did not modulate the recruitment of DLPFC, nor coupling between the hippocampus and VLPFC, across trial types is consistent with an age-related impairment in the ability to engage controlled retrieval and monitoring processes in accordance with task demands (Giovanello & Schacter, 2011; McDonough et al., 2013). A failure to engage such strategic retrieval processes (i.e. recall-to-reject) may have contributed to a reduction in correct rejection rate in the older group in the present study. More broadly, impaired engagement of such processes may contribute to age-related increases in false recognition, particularly under conditions in which one must disqualify experimentally family test cues as having been studied by engaging in strategic retrieval and monitoring of target details.

Unlike rejecting recombined pairs, which places considerable demand on strategic retrieval processes (Rotello & Heit, 2000; Lepage et al., 2003; Cohn & Moscovitch, 2007), endorsing intact pairs is thought to reduce these demands by facilitating access to stored details and minimizing demands on post-retrieval monitoring and evaluation processes. Here we tested the hypothesis that the retrieval support offered by intact pairs would increase the likelihood that associative hits would be supported by common neural mechanisms across age groups. In support of this possibility, the results of the univariate activity analyses revealed that both older and younger adults preferentially recruited the hippocampus and angular gyrus during hits relative to correct rejections. Moreover, the whole-brain exploratory results provided evidence that these areas were recruited in conjunction with other regions included in the core recollection network, including the medial prefrontal cortex and the posterior cingulate cortex, as well as areas associated with the retrieval of visual details, including the lingual gyrus and middle occipital cortex. The engagement of this set of brain areas when endorsing studied items has been well-documented in healthy younger adults (Kim, 2010; Rugg & Vilberg, 2013), whereas in older adults the evidence for such engagement has been mixed (Dennis, Kim, & Cabeza, 2008; Duarte, Graham, & Henson, 2010; de Chastelaine, Mattson, Wang, Donley, & Rugg, 2016). The present findings of age-invariant increases in the recruitment of these regions during hits as compared to correct rejections provides some evidence for the possibility that the presence of age-related differences in retrieval mechanisms may be determined by demands on controlled processes at retrieval. That is, under conditions in which greater retrieval support is provided and demands on controlled processes are reduced (i.e. in the form of a strong retrieval cue that minimizes retrieval and monitoring demands), age differences in the neural mechanisms supporting memory retrieval are less likely to be observed.

Notably, the engagement of regions within the core recollection network, and in particular the hippocampus and angular gyrus, have reliably been elicited during successful recollection of previous events, but not when recognition is based on a feeling of familiarity (Eldridge et al., 2000; Yonelinas et al., 2005; Wagner et al., 2005; Vilberg & Rugg, 2008). The age-invariant increase in recruitment of these brain regions during hits relative to correct rejections in the present study raises the possibility that older and younger adults were engaging similar mechanisms to endorse intact pairs, and that this mechanism involved recollection of the target associate. This type of strategy may have been facilitated by the correspondence between the stored representation and the retrieval cue (in this case, most likely based on semantic rather than perceptual information due to the absence of the pictorial stimuli at test). Such a possibility would be consistent with some existing behavioural observations of increased ability to access target details when endorsing intact pairs, as compared to rejecting recombined pairs (Cohn et al., 2008; Trelle et al., 2017). Future studies that employ more direct measures of the retrieval of target details, such as the explicit report of target details from encoding not provided in the test cue, the use of remember/know judgments at test, or multivariate metrics of cortical reinstatement of target features during retrieval, are required to provide more direct evidence for this possibility.

It is important to note that although the neural data provide evidence in favor of a selective age-related deficit in processes necessary to reject recombined pairs, or recall-to-reject, coupled with intact mechanisms for supporting target recollection, behaviorally older adults exhibited reductions in hit rate and correct rejection rate that were comparable in size. This pattern differs from some previous findings, in which age-related impairments in hit rate is often smaller than that for the correct rejection rate of non-studied lures. One factor that may have contributed to the equivalent impairment in the present study is the removal of the studied images from the test cues, which reduced the potential contribution of perceptual fluency to performance and encouraged the use of a recollection-based retrieval strategy across trial types. Importantly, perceptual fluency and familiarity-based retrieval processes are thought to contribute to the absence or minimization of age-related differences in hit rate in typical recognition memory studies (Jennings & Jacoby, 1993; Koen & Yonelinas, 2016). Minimizing the contribution of such processes in the present study, as was intended by the removal of studied images from the test phase, may have contributed to a comparable deficit in hits and correct rejections here. Although hits were indeed less frequent in older relative to younger adults, the results of the present fMRI data analysis suggest a common mechanism supported hits across groups, which may have involved target recollection.

Taken together, the present data provide further evidence for at least two dissociable factors underlying age-related declines in episodic memory, together with new insights into the neural mechanisms underlying age-related decline in episodic memory, and in particular the ability to discriminate between events that share overlapping elements. Here we provide neural evidence for both declines in the differentiation of stimulus representations during encoding, as well as reductions in the recruitment of controlled retrieval processes to reject recombined pairs, coupled with age-invariant mechanisms for endorsing intact pairs. These results complement our previous behavioural data (Trelle et al., 2017) in suggesting a contribution of age-related changes in representational quality and strategic retrieval processes to episodic memory decline with age. Moreover, the present findings help to explain disproportionate deficits in recall-to-reject relative to recall-to-accept processes among older adults, and suggest that age-related declines in recollection-based retrieval processes vary as a function of demands on controlled retrieval processes.

## Acknowledgements

We would like to thank the staff of the MRC Cognition and Brain Sciences Unit MRI facility for scanning assistance and Deborah Green for assistance with data collection.

