## Supplementary Materials for "Neural evidence for age-related differences in representational quality and strategic retrieval processes"

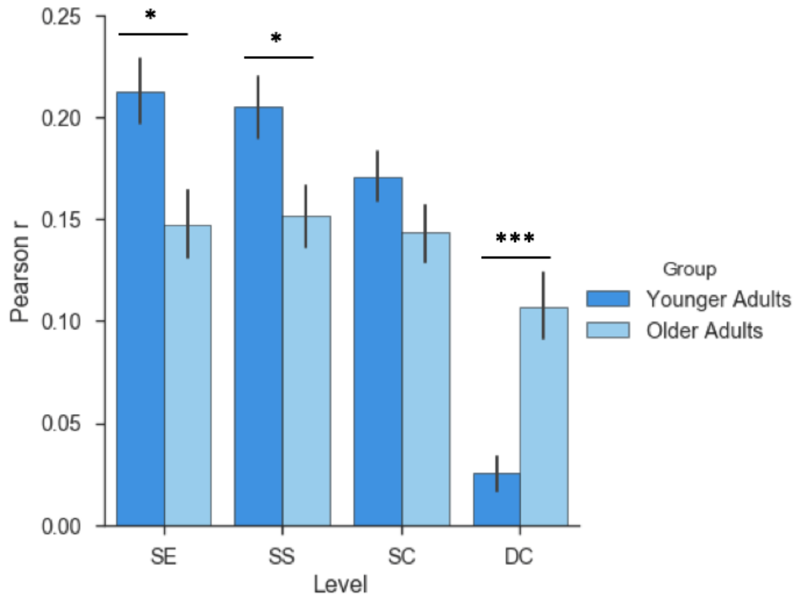


*Figure S1: Pattern similarity at encoding for subsequently remembered stimuli as a function of event relatedness. Relative to younger adults, older adults exhibit reduced pattern similarity for related events, coupled with increased pattern similarity for events that are more distinct. Error bars represent standard error of the mean.* ***$p < .001$; * $p < .05$

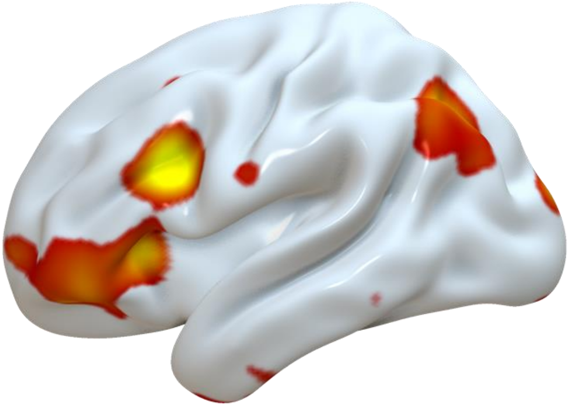

*Figure S2: Univariate Age x Condition (Hits > CRs) interaction. Uncorrected whole brain maps thresholded at p* < .005.

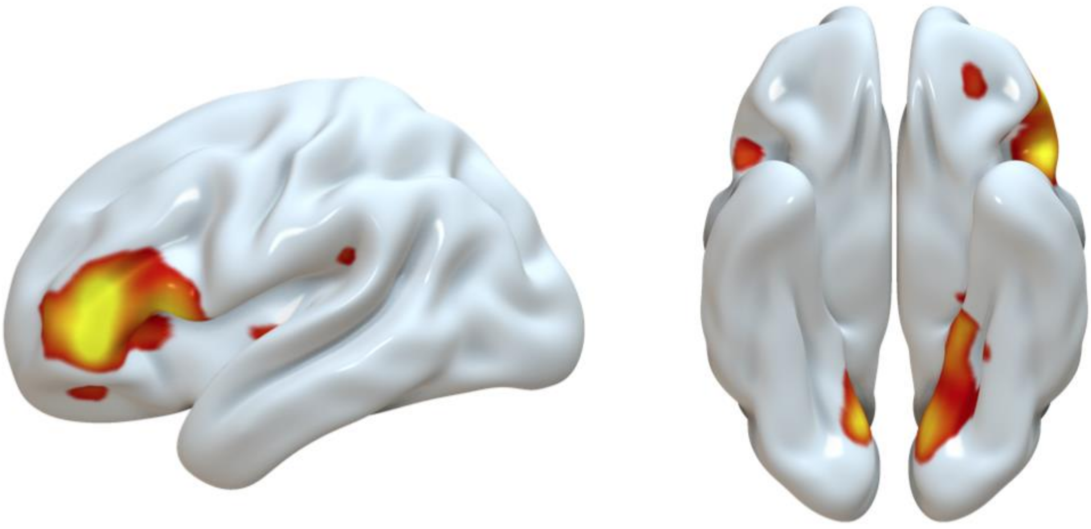

*Figure S3: Hippocampal connectivity Age x Condition (Hits > CRs) interaction. Uncorrected whole brain maps thresholded at p* < .005.

*Table S1: Regional BOLD activity during Hits and CRs. Uncorrected whole brain results thresholded at p* < .005. *Asterisk (*) indicates effects that survive FWE p* < .05

| Region | Voxels | MNI Coordinates (x, y, z) | | | Peak *t* value |
|---|---|---|---|---|---|
| ***Hits > CRs*** | | | | | |
| **L hippocampus*** | 468 | -27 | -30 | -12 | 7.96 |
| **L superior medial frontal gyrus*** | 1262 | -6 | 51 | 6 | 7.65 |
| **L lingual gyrus*** | 282 | -12 | -78 | -9 | 7.51 |
| **L inferior parietal cortex*** | 351 | -54 | -45 | 39 | 7.37 |
| **R cuneus*** | 163 | 12 | -93 | 21 | 7.24 |
| **R middle occipital gyrus*** | 71 | 42 | -81 | 24 | 6.82 |
| **R hippocampus*** | 186 | 27 | -18 | -18 | 6.76 |
| **L middle cingulate cortex*** | 393 | 0 | -18 | 39 | 6.33 |
| **R supramarginal gyrus** | 147 | 57 | -30 | 45 | 6.28 |
| **R insula** | 37 | 27 | 18 | -15 | 5.87 |
| **L inferior temporal gyrus** | 15 | -54 | -9 | -27 | 5.01 |
| **R middle temporal gyrus** | 15 | 54 | -30 | -6 | 4.81 |
| **L middle temporal gyrus** | 20 | 57 | -51 | 3 | 4.64 |
| **L insula** | 19 | -36 | -6 | 18 | 4.63 |
| ***CRs > Hits*** | | | | | |
| **R lingual gyrus*** | 160 | 18 | -75 | -9 | 7.11 |
| **L cuneus** | 56 | -6 | -93 | 9 | 5.00 |
| **L superior frontal gyrus** | 52 | -24 | -9 | 60 | 4.68 |
| ***Group X Condition*** | | | | | |
| **L inferior frontal gyrus (triangularis)** | 106 | -36 | 21 | 27 | 3.55 |
| **L inferior parietal lobe** | 25 | -33 | -69 | 39 | 3.17 |
| **L insula** | 19 | -30 | 27 | 3 | 3.11 |
| **L precuneus** | 19 | -6 | -48 | 12 | 2.96 |

*Table S2: Hippocampal Connectivity during Hits and Correct Rejections. Uncorrected whole brain results thresholded at p* < .005.

| Region | Voxels | MNI Coordinates (x, y, z) | | | Peak *t* value |
|---|---|---|---|---|---|
| ***Hits > CRs*** | | | | | |
| *No suprathreshold clusters* | | | | | |
| ***CRs > Hits*** | | | | | |
| **L pallidum** | 48 | -9 | 6 | -3 | 4.47 |
| **R temporal Pole** | 85 | 54 | 6 | -12 | 4.16 |
| **L temporal Pole** | 36 | -48 | 15 | -24 | 4.14 |
| **L amygdala** | 15 | -15 | -6 | -18 | 3.94 |
| **R caudate** | 16 | 18 | -9 | 24 | 3.66 |
| **R inferior frontal gyrus (orbitalis)** | 17 | 48 | 27 | -6 | 3.36 |
| ***Group X Condition*** | | | | | |
| **L inferior frontal gyrus (triangularis)** | 306 | -45 | -42 | 12 | 3.84 |
| **L fusiform gyrus** | 28 | -21 | -39 | -12 | 3.58 |
| **L superior occipital gyrus** | 12 | -15 | -90 | 12 | 3.39 |
| **L inferior occipital gyrus** | 24 | 30 | -87 | -6 | 3.37 |
